# Medial entorhinal spike clusters carry more finely tuned spatial information than single spikes

**DOI:** 10.1101/2020.06.10.144204

**Authors:** Leo. Richard Quinlan, Susan. Margot Tyree, Robert. Gordon. Keith. Munn

**Affiliations:** Discipline of Pharmacology, National University of Ireland, Galway; Discipline of Physiology, National University of Ireland, Galway; Department of Neurobiology, Stanford University School of Medicine

## Abstract

Many cells within the entorhinal cortex (EC) fire relatively infrequently, with the majority of their spikes separated by many hundreds of milliseconds. However, most cells are seen to occasionally fire two, three, or more spikes in quick succession. Recent evidence has shown that, in EC grid cells, “burstier” cells; cells that fire more of their spikes in bursts, have more well defined spatial characteristics than cells that fire fewer bursts. However, there is evidence that the window for considering related spikes in MEC could be as long as 100ms. Here, we divide the spikes fired by single cells into single spikes and “clusters” of spikes occuring within 100ms. We show that these burst “clusters” of spikes fired by cells in MEC convey more finely tuned spatial and directional information than the numerically more common single spikes. In addition, we find that introducing environmental uncertainty decreases the ratio of clusters fired to single spikes. Most crucially, we find that although single spikes are less spatially precise than clusters, they are more temporally precise – these spikes are more closely entrained to LFP theta than clusters. These findings demonstrate that clusters of spikes in EC convey more specific information about space than single spikes, may reflect “certainty” about spatial position and direction, and may represent a different firing “mode” in which intraregional communication is less relevant than interregional traffic.

## Introduction

The entorhinal cortex (EC) forms a central hub in the neural network that supports memory formation and navigational guidance in all mammals. The EC is part of the medial temporal lobe, and constitutes a major connection between sensory and parahippocampal cortices and the hippocampus (HPC)^1,2^. As a sensory convergence area, EC represents the first step in the integration of sensory information into a coherent spatial and temporal representation. The medial portion of the entorhinal cortex (MEC) is rich in diverse, functionally well-defined cell types. These cells include the regularly tessellating firing fields of grid cells^3^, the directionally specific firing of head direction cells^4,5^, and the speed-correlated firing of speed cells^6,7^. While these definitions are classically defined on the basis of their firing-rate tuning curves, recent advances in computational and statistical modelling have shown that cells in MEC are much more plastic and heterogenous in their coding properties than previously thought^8^. Many of the neurons in this region code for several different spatial variables simultaneously, and there is no doubt that many other types of information are encoded here also^9,10^. The emerging view of MEC is that of a dynamic and flexible region in which multiple variables can be coded simultaneously by single neurons.

Most of the principal neurons in MEC have relatively low firing rates, only occasionally firing trains of spikes longer than single spikes. However, there is evidence that these trains although rare are different from isolated spikes; recent data has suggested cells in MEC can be classified into “bursty” and “non-bursty” groups through analysis of their interspike interval (ISI) distributions^11^. Burst coding is thought to be computationally advantageous compared to single spiking^12–14^. It has been established that the propensity of cells to fire bursts within hippocampus and subiculum is related to increased spatial selectivity^15,16^. More recently, in MEC it has been shown that grid cells that fire more of their spikes as bursts relay more precise spatial information than grid cells that burst-fire less often^17^. It has been suggested that burst coding relays a distinct stream of information from single spikes^18^. These lines of evidence lead to the suggestion that there is a clear delineation between isolated spikes and spikes that occur as bursts within short timescales from one another. One possible way in which these ensembles might be organized and segregated from one another is their activity relative to the local field potential (LFP).

Theta rhythm (6-12 Hz) in the hippocampus and MEC is thought to coordinate the firing of the neurons in these regions, segregating the input/output flow of information between them ^19,20^. Theta rhythm in hippocampus and MEC is conducted largely from the medial septum (MS), which sends strong cholinergic afferents to both MEC and HPC^21^. The cholinergic projections to MEC terminate in layers II and V, which are the layers of MEC that project to HPC^22^, further suggesting a role for MS cholinergic innervation in coordinating information between MEC and HPC. Critically, muscarinic receptor activation has been shown to coordinate the firing of neurons *in vitro*, and induces coordinated discharges remarkably reminiscent of spontaneous spike bursts^23^. It is possible that the coordinated spiking of neurons in MEC *in vivo* is organised through cholinergic projections from MS, and that this coordinated activity is more likely to result in burst firing rather than single spike generation.

Apart from the traditional classification of spikes firing as bursts (occurring within 10 ms of each other), there is reason to believe that cells in MEC and HPC that repeat fire within a theta cycle (∼100 ms) may be associated in a similar way to bursts. *In vitro* stimulation at theta frequencies (i.e. 100 ms spaced intervals) is more effective at potentiating synapses than high frequency stimulation^24^, suggesting that these cells are intrinsically tuned to respond preferentially to theta-scale inputs. Indeed, intrinsic calcium dynamics have been shown to transition granule cells in cerebellum between single spiking and bursting modes^25^. One of the dominant types of intrinsic oscillation observed in MEC is driven by the hyperpolarization-activated cyclic nucleotide–gated (HCN) channel inward current (I_h_). This oscillation occurs at roughly theta frequency and regulates the frequency of septal-conducted theta^26–28^. Suggesting that spikes fired within the timescale of a theta oscillation might be computationally associated. Indeed, it has been noted that MEC neurons have a tendency to fire temporally related spikes on the timescale of I_h_. These related spikes have been dubbed “clusters”^29^.

Determining coding in behaviorally correlated cell types such as grid and head direction cells requires animals to behave, but moreover it requires the cell to fire a number of spikes during that behavior that allows spiking patterns to be meaningfully analyzed. Analyzing bursts in isolation from single spikes is therefore challenging as cells in MEC typically fire relatively few classically-defined bursts This ratio of spikes to bursts makes analysis problematic, as differences in behavioral sampling can differ substantially between groups. To circumvent this issue, we use a more liberal classification of bursts previously defined as “clusters” as in Alonso and Klink^29^. This division facilitates the capture of spikes traditionally classified as bursts (i.e. within 10 ms of one another) as well as other spikes that are nonetheless likely computationally associated, since they occur on the timescale of a theta oscillation, and also on the timescale of I_h_. The larger capture of spikes using this definition allows analysis of clusters of spikes in isolation from single spikes in a way not possible using more restrictive criteria.

Here we examine an existing dataset of many hundreds of MEC neurons ^30^ to determine whether cluster spiking in MEC is quantitatively different from single spike events. Using a large population of cells, we first show that the traditional bursting characteristics of MEC are similar to those previously reported^11^, namely that there is a clear delineation between “bursty” and “non-bursty” cells. We then dissect the spiketrains of MEC cells into single spikes and the more liberally defined “clusters” of spikes that allows us to examine these separate groups of spikes in isolation from each other. We find that the tuning curves of all cell types in the MEC are sharper and more stable when only the clusters are considered, while a less precise and more variable code is carried by the single spikes. Moreover, introducing spatial uncertainty causes the proportion of spikes fired as clusters to decline markedly, suggesting that clusters of spikes are associated with well-known or certain information. Lastly, we show that single spikes are more phasically locked to LFP theta than clusters, suggesting different functional properties of these two firing modes. These findings have important implications for our understanding of the neural coding of space within the MEC, and the coding strategies of neurons more generally.

## Results

Wildtype mice (n = 25 mice) explored a familiar square environment while recording to screen for cells of interest. On recordings in which it was determined that a cell was recorded, the animals were also recorded in an unfamiliar rectangle. A total of n = 193 cells were recorded. Of these 16 % (n = 31 cells) were identified as grid cells, 18.6 % (n = 36 cells) as head direction cells, and 27.5 % (n = 53 cells) as speed cells.

### Cells recorded from MEC fire traditional bursts of spikes and can be classified into two groups

Previous studies have shown that cells in the MEC can be classified into “bursty” cells that fire many spikes with an interspike interval of less than 10 ms, and non-bursty cells that fire the majority (> 50 %) of their spikes with interspike intervals longer than 10 ms (Figure 1D, Ebbesen et al., 2016). In our current dataset, we also observe this range of activity (Figure 1A,B). Principal component analysis (PCA) of the ISI histograms of each cell demonstrated the classic ‘C’ shape of the projection of the first two principal components (Figure 1E). The majority of variance in the dataset was captured by the first three principal components with the first two alone capturing nearly half of the variance. K-means clustering robustly separated the cells into two clusters based on their principal components (Figure 1F), and this clustering closely agreed with classification of cells based on burst score: their ratio of “burst” spikes (with ISIs less than 10 ms) against the total number of spikes (Figure 1C) (i.e mean time between spikes)

**Figure 1.**
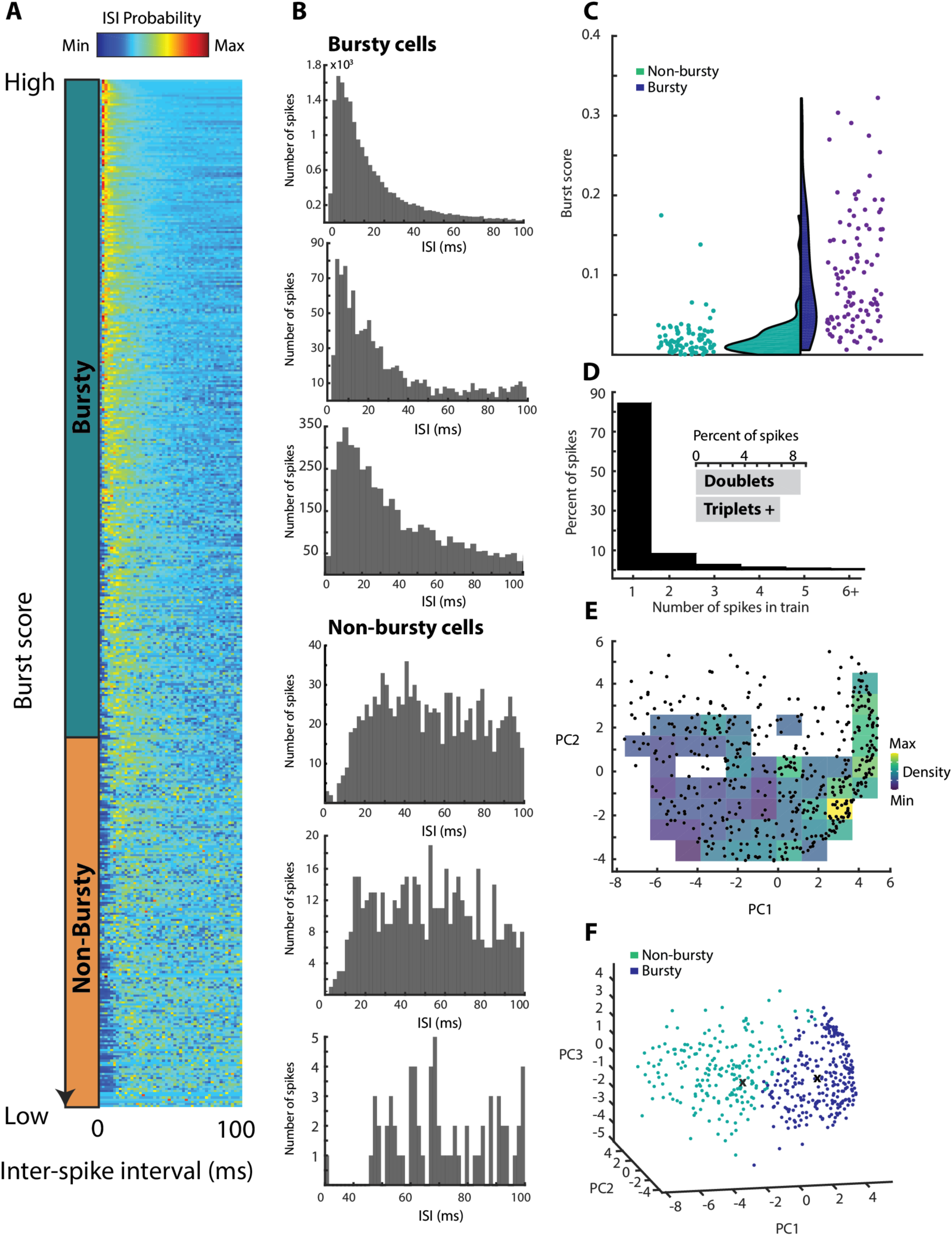
The majority of MEC neurons are classified as “bursty”. **A)** Each row (n = 475 neurons) represents the probability of a cell firing in a particular isi bin (2 ms bins) from 0 to 100 ms. Cells are sorted by their burst score (defined as the number of spikes with an isi < 10 ms divided by the total number of spikes in 100 ms). A bar to the left of the figure shows cells classified as bursty (green) and non-bursty (orange) through k-means clustering of the principal components of the ISI distributions **B)** Histograms illustrating the ISI distributions of six individual neurons; three classified as bursty (top three) and non-bursty (bottom three). Bursty cells fired the majority of their spikes from 0-100 ms within 20 ms of the previous spike, while the majority of spikes fired by non-bursty cells from 0-100 ms were more than 20 ms from the previous spike. **C)** The burst score (spikes with ISI < 10 ms/ total spikes in 100 ms) of cells classified as bursty are generally higher than cells classified as non-bursty by k-means clustering of the principal components of the ISI distributions. Half-violins demonstrate the density of the distributions of values. **D)** The percentage of all spikes fired across all neurons as single spikes or in bursts of two or more spikes. The overwhelming majority of all spikes were fired as single spikes in any 10 ms time window (84.56%). The inset compares the percentage of spikes fired as doublets compared to all of the spikes fired in trains of three or more spikes. The majority of burst spikes were doublets (8.56%), compared to 6.88% of spikes that occurred in bursts of three or more. **E)**. Projection of the first three principal components of the ISI histograms for each cell. A K-means clustering algorithm clearly separates cells into “bursty” and “non-bursty” groups as in Ebbesen et. al, 2016. The centroids of each cluster are marked with an ‘x’). **F)** Principal components analysis of the ISI histograms of each neuron, projected onto the first and second principal components. Colored background shows the relative density of neurons in PC space. As in Latuske et. al, (2015) and Ebbesen et. al, (2016), the ISI histograms make a c-shaped projection onto PC space for the first two principal components.

### There are systematic differences in the morphology of spikes fired in clusters compared to spikes fired singly

The waveforms of spikes identified as belonging to a cluster were compared to spikes fired singly. Neurons that fired high amplitude single spikes also had high amplitude cluster spikes and vice-versa (r = 0.927, p = 1.60^-83^). However, comparing within neurons, spikes fired as singletons had an average higher peak/trough amplitude (Figure 2A-C, Table 1) than clustered spikes. Cluster spikes were also narrower than single spikes (Figure 2A-B,2D, Table 1), although the time from peak to trough did not differ between them (Figure 2A-B,2E, Table 1). Critically, this difference in amplitude was primarily driven by the fact that single spikes originated from a more hyperpolarized initial potential than clusters, which began when the cell was already somewhat depolarized (mean initial potential ± SEM (mV), single spikes = -1.87 ± 0.45, clusters = -1.079 ± 0.30, Z = 3.43, p = 6.11^-4^). When the absolute maximum height of the spike was considered, clusters reached higher amplitudes than single spikes (mean max amplitude ± SEM (mV), single spikes = 45.20 ± 0.80, clusters = 46.42 ± 0.63, Z = 5.22, p = 1.78^-7^). Single spikes also had deeper afterhyperpolarizations, reaching lower potentials than cluster spikes (mean minimum trough amplitude ± SEM (mV), single spikes = -16.62 ± 0.65, clusters = -13.80 ± 0.47, Z = 9.25, p = 2.16^-20^).

**Table 1.** Characteristics of spikes fired as clusters compared to spikes fired as singletons (means ± SEM)

| Feature | Single Spikes | Clusters | Statistic |
| --- | --- | --- | --- |
| Amplitude<br>(peak/trough, μV) | $61.83 \pm 1.27$ | $60.22 \pm 0.91$ | $Z = 2.74$, $p = 0.006$ |
| Spike width (s) | $0.130 \pm 0.003$ | $0.123 \pm 0.002$ | $Z = 3.18$, $p = 0.002$ |
| Spike latency (s) | $0.386 \pm 0.013$ | $0.370 \pm 0.011$ | $Z = 1.01$, $p = 0.314$ |

**Figure 2.**
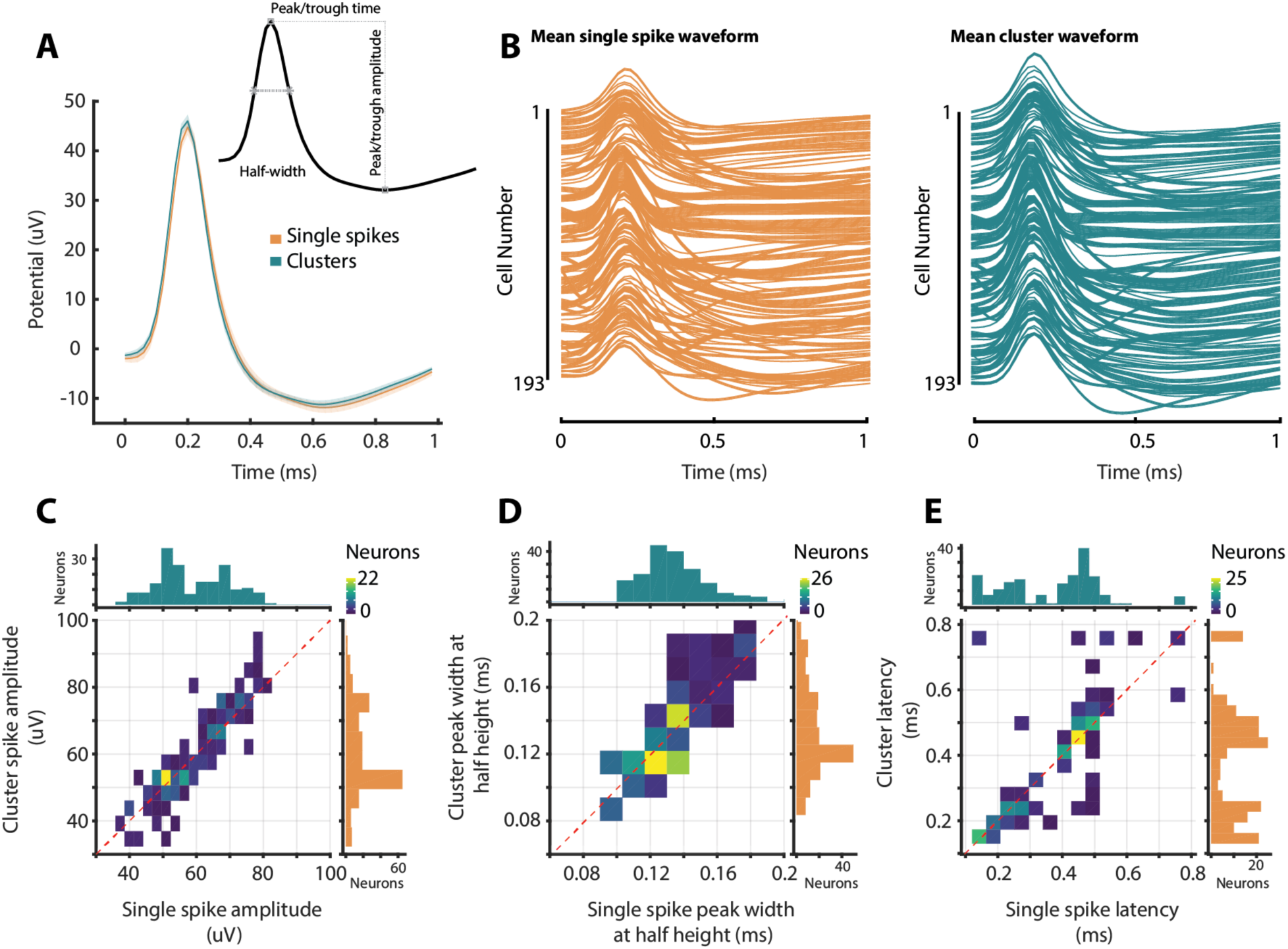
Characteristics of spikes fired as part of a cluster differ from those fired singly. **A)** The mean waveform over all spikes fired by all 193 neurons in clusters (green) and singly (orange). The shaded region indicates the 95% CI. Clusters had wider and higher amplitude peaks than spikes fired outside of clusters. Inset demonstrates the points at which spike measurements were made. **B)** The mean waveform for the single spikes (left, orange) and clusters (right, green) of each cell. Cells are stacked vertically according to their order of recording. **C-E)** Density plots of (A) peak-to-trough amplitudes (B) Half-width, and (C) peak/trough latency of spikes fired as singletons and spikes fired as clusters by the same neuron. Marginal histograms illustrate the distributions of peak to trough amplitudes of single spikes (orange) and clusters (green). Red dotted lines illustrate unity between these variables.

### Clusters of spikes by grid cells are much more spatially well-defined than single spikes

Grid cells were defined based on shuffled distributions as previously described ^30^ (n = 14 mice, n = 31 grid cells). The spikes fired by these grid cells were divided into single spikes and clusters, the longer spiketrain was always downsampled to match as described in the methods. Unlike most cells, grid cells often fired more than 50 % of their spikes in clusters, leading to the cluster spiketrains being downsampled. As can be seen from the rate maps generated by the clusters compared to the single spikes for two cells (Figure 3A), the apparent nodes of the grid pattern, although similar between clusters and single spikes, are better defined and have a higher contrast between field and ground when examining the clusters in isolation from the single spikes. The cluster firing of grid cells carries more spatial information (Figure 3B, mean information ± SEM (bits/spike), single spikes = 0.372 ± 0.088; clusters = 1.042 ± 0.049, Z = 4.80, p = 1.578^-6^), and are more spatially precise (Figure 3C, mean spatial selectivity ± SEM, single spikes = 5.87 ± 0.807; clusters = 24.76 ± 4.28, Z = 4.86, p = 1.174^-6^) and compact (Figure 3D, spatial sparsity ± SEM, single spikes = 0.661 ± 0.031; clusters = 0.328 ± 0.029, Z = 4.86, p = 1.174^-6^) than the single spikes of these cells. In agreement with previous observations indicating increased spatial tuning when animals run at high velocities ^8^, we find that clusters of spikes are more likely than single spikes to occur when the animal is running at high speed (Figure 3E).

**Figure 3.**
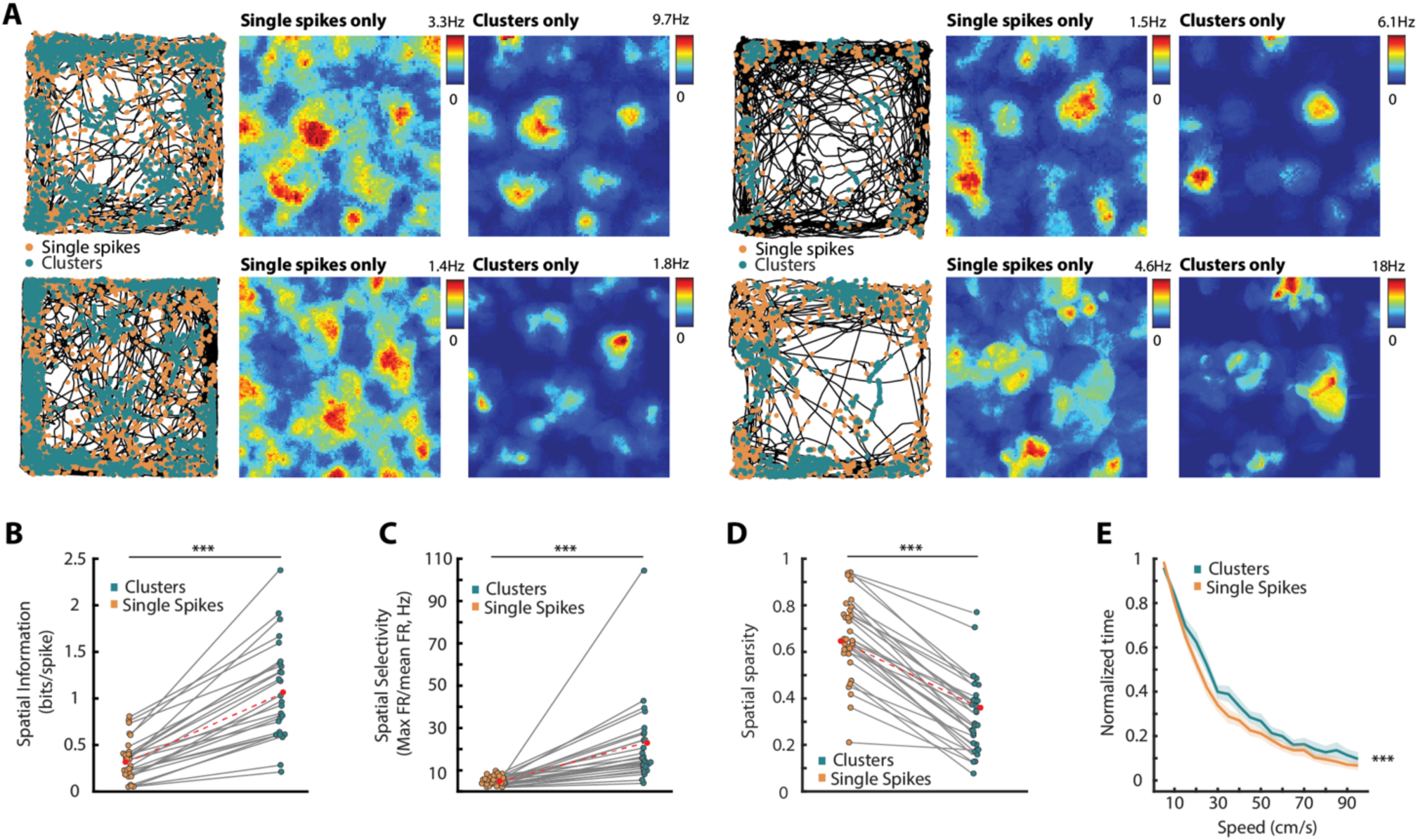
Grid Cell clusters are more spatially precise than single spikes. **A)** Two example grid cells; Left, the path taken by an animal over the course of a recording session (black line) overlaid with the spatial positions that each spike fired. Spikes are divided into single spikes (orange) and cluster spikes (green). Cluster spikes are largely confined to the nodes of each grid cell compared to single spikes which are more spatially distributed. Center) Adaptively smoothed rate map generated by only the single spikes of each cell. Right) Adaptively smoothed rate map generated by only the clusters of each cell. Rate maps generated from clusters are generally better defined than those generated from single spikes **B)** The information content of clusters of spikes (green) was substantially greater than the information contained by single spikes (orange). Red dots indicate the means of each group of spikes, and the difference between them is illustrated with a dashed red line. **C)** There was a greater difference between the mean and maximum firing rates when considering only the cluster spikes compared to single spikes. **D)** Clusters of spikes fired by grid cells were significantly less spatially sparse than single spikes **E)** Spikes that were fired in clusters occurred more often when the animal was running faster than when spikes were fired as singlets (mean speed ± SEM (cm/s), Single spikes = 27.16 ± 1.20; Bursts = 29.78 ± 1.38, Z = 3.67, p < 0.001). All panels: means are indicated with a red point connected by a red dashed line. Grey lines connect the single spikes and clusters fired by a single cell. ***p < 0.001

### Clusters of spikes carry more directional information than single spikes

As a population, MEC cells (n = 193 cells) coded direction relatively poorly. In spite of this poor coding of direction, nevertheless, the clusters of spikes fired by the population were significantly more directional than the equivalently downsampled number of single spikes (Mean directional vector length ± SEM, single spikes = 0.173 ± 0.013; clusters = 0.231 ± 0.015, Z = 7.322, p = 2.45^-13^, Figure 4B). As might be expected, this difference was largely driven by cells that were defined as “directional” (vector length over 0.2, Figure 4A) using traditional shuffling methods. When considering only these neurons (n = 12 mice, n = 37 directional cells), cluster spikes were observed to carry much more finely tuned directional information in comparison to single spikes (mean vector length, cluster spikes = 0.348 ± 0.035; single spikes = 0.227 ± 0.021; Z = 4.115, p = 3.867×10^−5^ Figure 4C). Likewise, the directional information content (as in Baird et al., 2001) of cluster spikes was much greater than the information relayed by single spikes (Figure 4D, directional information ± SEM (bits/spike), single spikes = 0.081 ± 0.011; clusters = 0.135 ± 0.018, Z = 5.016, p = 5.27^-7^). Head direction cells were equally directionally stable whether considering their single spikes or clusters (Figure 1E, directional stability ± SEM, single spikes = 0.754 ± 0.039; clusters = 0.796 ± 0.033, Z = 1.037, p = 0.300, n.s). Indicating that the low directional vector of the single spikes was not due to directional instability.

**Figure 4.**
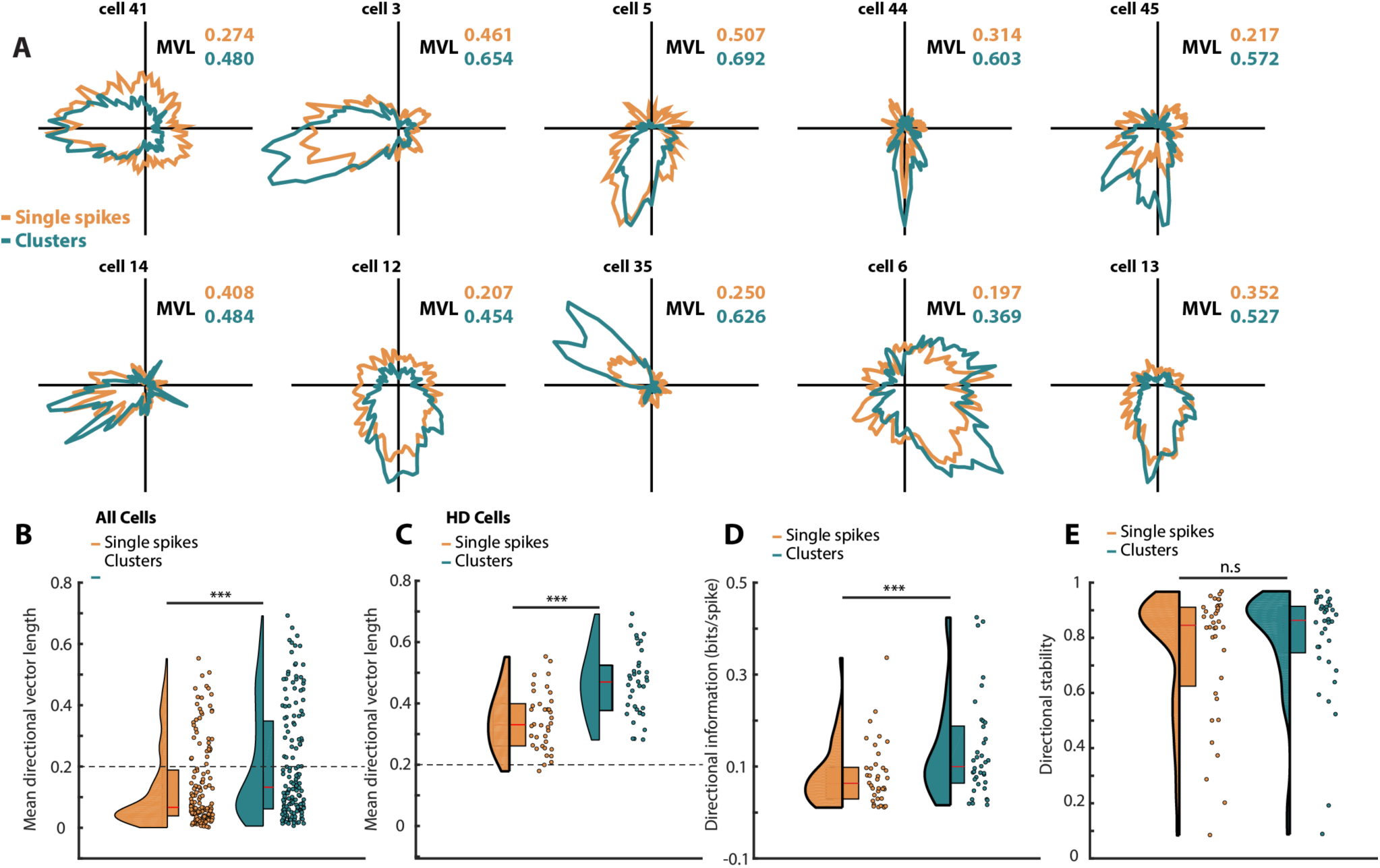
Clusters fired by MEC head direction cells are more finely tuned than single spikes. **A)** Firing rate tuning curves of ten head direction cells. Tuning curves are separated into that generated from single spikes (orange) and clusters of spikes (green). Cluster-derived tuning curves tended to be much more precise than the broader curves generated from single spikes. Alongside each turning rate curve pair for each cell is the color-coded mean directional vector length (MVL) of each tuning curve. **B)** As a population, MEC cells were relatively directionally non-specific. Nevertheless, clusters of spikes (green) fired by these cells were more directionally specific than single spikes (orange). A black dashed line indicates the vector length threshold at which cells are considered to be “directional”. **C)** When considering only cells whose full spiketrain passes the threshold to be considered directional, single spikes, although directionally specific enough to be considered “directional” in all but one case, are significantly less directionally specific than the clusters fired by these same cells. As in (B), a dashed line indicates the threshold at which cells are considered directional. **D)** In addition to being more directionally specific, clusters conveyed a greater amount of directional information than single spikes. **E)** While clusters were more directionally specific and informative, they were no more directionally stable than single spikes. In panels B-E, half violins indicate the density of the underlying distribution of values, while half boxplots show the first and third quartile of the distributions, while the median of each group is illustrated on the boxplot with a solid red line. *** p < 0.001, n.s = not significant

### Clusters relay more, and more temporally stable, information about speed than single spikes

Cells were defined as speed cells by reaching speed score and stability thresholds (as defined methods, below), and firing at least 100 spikes in bursts (n = 17 mice, n = 53 speed cells). These criteria were computed from their complete spiketrains. Clusters fired by speed cells produced a firing rate that was more tightly correlated with running speed than their respective downsampled single spikes (Figure 5A,E, Speed/firing rate correlation, ± SEM, single spikes = 0.105 ± 0.011; clusters = 0.133 ± 0.15, Z = 2.147, p = 0.0318). This correlation was also more stable over an entire recording session for clusters compared to single spikes (Figure 5B, Speed/firing rate correlation stability ± SEM, single spikes = 0.416 ± 0.041; clusters = 0.524 ± 0.037, Z = 2.023, p = 0.0431). Moreover, the slope and intercept of the speed/firing rate relationship of these cells were both higher when considering clusters compared to single spikes (Figure 5C,D, Slope (Hz/(cm/s)) ± SEM, single spikes = 0.0070 ± 0.003; clusters = 0.0092 ± 0.002, Z = 5.236, p = 1.637 x 10^−7^. Intercept (Hz) ± SEM, single spikes = 0.261 ± 0.076; clusters = 0.340 ± 0.088, Z = 3.191, p = 0.0014).

**Figure 5.**
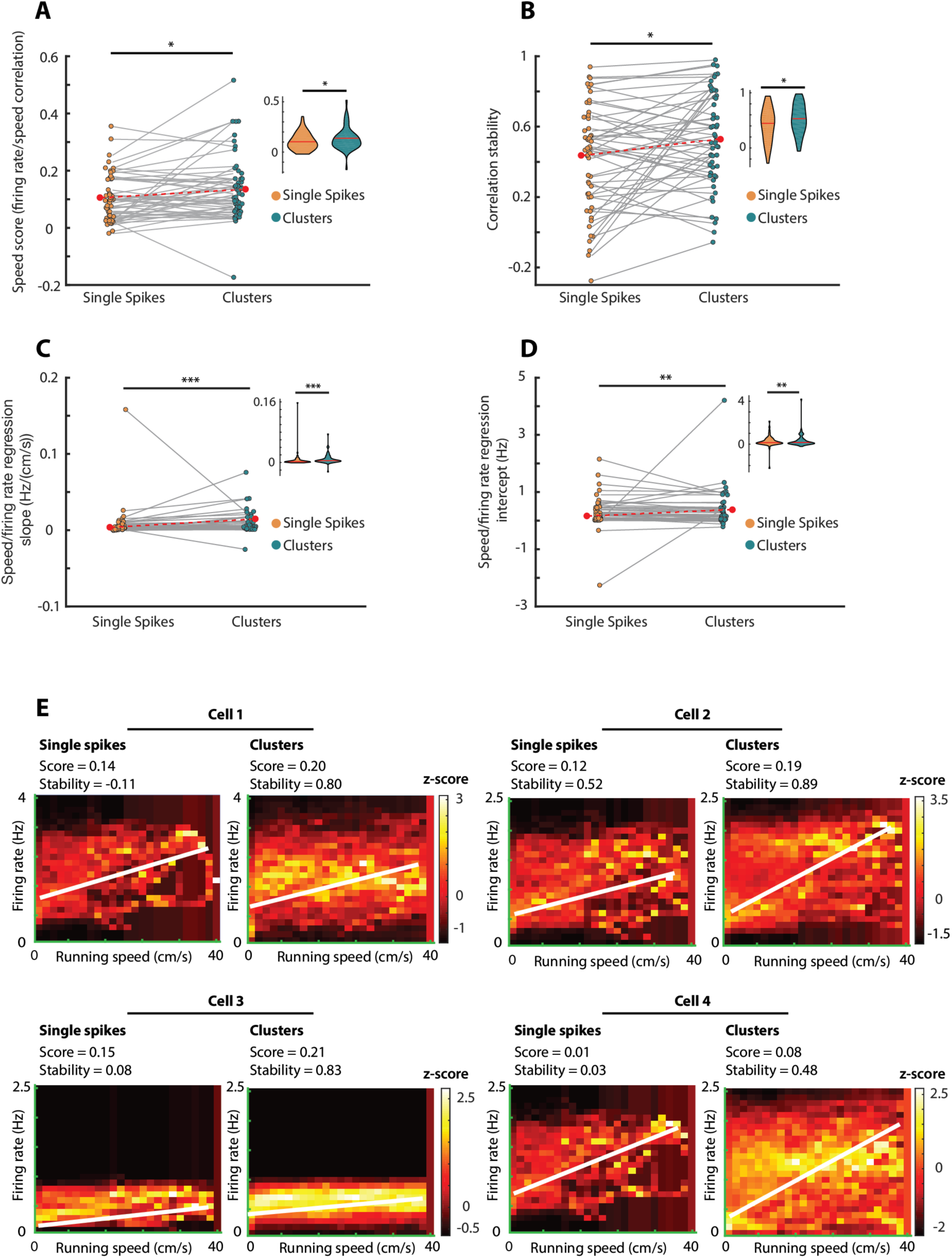
Clusters convey richer speed-related information than single spikes. **A)** The speed score derived from correlation of the firing rate with running speed is higher for clusters of spikes (green) compared to single spikes downsampled to numerical equivalence (orange). Grey lines connect the speed score produced from the single spikes with the score produced from the clusters of each cell. Inset: Violin plot illustrating the distributions of values, mean illustrated with a solid red line **B)** The correlation between firing rate and running speed was more stable over the recording session for clusters of spikes compared to single spikes. Inset: distribution of stability values. **C), D)**, The slope (C) and intercept (D) of linear regression through the firing rate/running speed relationship was higher for clusters of spikes compared to single spikes. Insets: Distributions of values. In all paired plots, red dots illustrate the means, and are connected by a dashed red line to illustrate the difference in values between single spikes and clusters. In all violin plots of distributions red lines show the mean of the distributions. * p < 0.05, ** p < 0.01, *** p < 0.001 **E)** Z-scored firing-rate by running speed probability plots for four individual speed cells. The left side of each cell’s plots are the single spikes fired by that cell, while the right hand panel is the plot produced from the clusters fired by the cell. For each plot, firing rate is binned by running speed into speed increments of 2.5cm/s, and then z-scored for each 2.5 bin. Hotter colors indicate a greater probability that a cell had a certain firing rate when running at a specific speed. Each of the cells shows the speed score (correlation between firing rate and running speed) and split-quarter stability for both the isolated single spikes and isolated clusters.

### Cells fire fewer of their spikes in clusters in unfamiliar environments

If clusters of spikes serve the function of consolidating information, or representing strong or certain information about the position of an animal in an environment, clusters should occur with less regularity when the animal is uncertain about an environment. The same neurons were recorded in a familiar square environment and an unfamiliar rectangular environment to which animals had infrequent exposure (fewer than 10 times total over 3-10 weeks, Figure 6A). As hypothesised, the overall number of spikes fired in clusters was less in the unfamiliar rectangle compared to the familiar square (Figure 6B, mean number of spikes ± SEM; familiar square = 1,534 ± 168, unfamiliar rectangle = 795 ± 91. Z = 6.898, p = 5.27^-12^). Critically, the proportion of spikes fired in clusters relative to single spikes was also lower in the unfamiliar rectangle than the familiar square (Figure 6C, proportion of spikes fired in clusters ± SEM; familiar square = 0.314 ± 0.010, unfamiliar rectangle = 0.282 ± 0.011, Z = 2.722, p = 0.007), showing that the difference was due to a reduction in the tendency to fire clusters in the unfamiliar environment. Although cells fired fewer of their spikes as clusters in the unfamiliar environment, clusters of spikes fired by grid cells still carried dramatically more spatial information than single spikes in the unfamiliar environment (Figure 6D-F, mean spatial information (bits/spike) single spikes = 0.376 ± 0.090, clusters = 0947 ± 0.103. Z = 4.566, p = 4.971^-6^). Clusters of spikes fired by grid cells in the unfamiliar environment were also more spatially selective than single spikes (mean spatial selectivity SEM; single spikes = 21.52 ± 16.99, clusters = 27.71 ± 10.93. Z = 4.253, p = 2.114^-5^). However, there was no overall difference in the magnitude of change between environments in information, selectivity, or stability between single spikes and clusters (Figure 6E,F). For cells identified as directional that fired more than 100 spikes as clusters in both environments (n = 27), there was a general reduction in mean vector length (Figure 6G,H mean vector length ± SEM, familiar square = 0.416 ± 0.024, unfamiliar rectangle = 0.282 ± 0.030, Z = 5.300, p = 1.16^-7^), information content (information content (bits/spike) ± SEM, familiar square = 0.120 ± 0.020, unfamiliar rectangle = 0.079 ± 0.015, Z = 3.500, p = 4.65^-4^) and directional stability (directional stability ± SEM, familiar square = 0.795 ± 0.045, unfamiliar rectangle = 0.561 ± 0.051, Z = 4.526, p = 6.01^-6^) when in the unfamiliar rectangular environment compared to the familiar square environment. However, although vector length and information content appeared to reduce more between environments for cluster spikes there was no difference between them (Figure 6I, mean vector length difference ± SEM, single spikes = -0.118 ± 0.023, clusters = -0.149 ± 0.034, Z = 1.081, p = 0.300, n.s; information content difference (bits/spike) ± SEM, single spikes = -0.030 ± 0.011, clusters = -0.050 ± 0.022, Z = 0.460, p = 0.650, n.s). Likewise, although cluster spikes seemed to lose less of their directional stability than burst spikes, there was no difference between single spikes and clusters (Figure 6I, stability difference ± SEM, single spikes = -0.240 ± 0.060, clusters = -0.209 ± 0.059, Z = 0.648, p = 0.517, n.s). As with directional cells, speed cells had lower speed scores and speed/firing rate correlation stability in the novel rectangular environment compared to the familiar square (Figure 6J, speed score, familiar square = 0.120 ± 0.012; novel rectangle = 0.053 ± 0.016, Z = 4.430, p = 9.425^-6^. Stability, familiar square = 0.470 ± 0.034; novel rectangle = 0.220 ± 0.031, Z = 5.328, p = 9.955^-8^). However, single spikes lost an equivalent amount of speed information between environments as clusters (Figure 6K; difference in speed score, single spikes = -0.0831 ± 0.027; clusters = -0.0513 ± 0.0130, Z = 1.658, p = 0.0974, n.s. Difference in stability, single spikes = -0.266 ± 0.043; clusters = -0.255 ± 0.042, Z = 0.610, p = 0.5418, n.s.).

**Figure 6.**
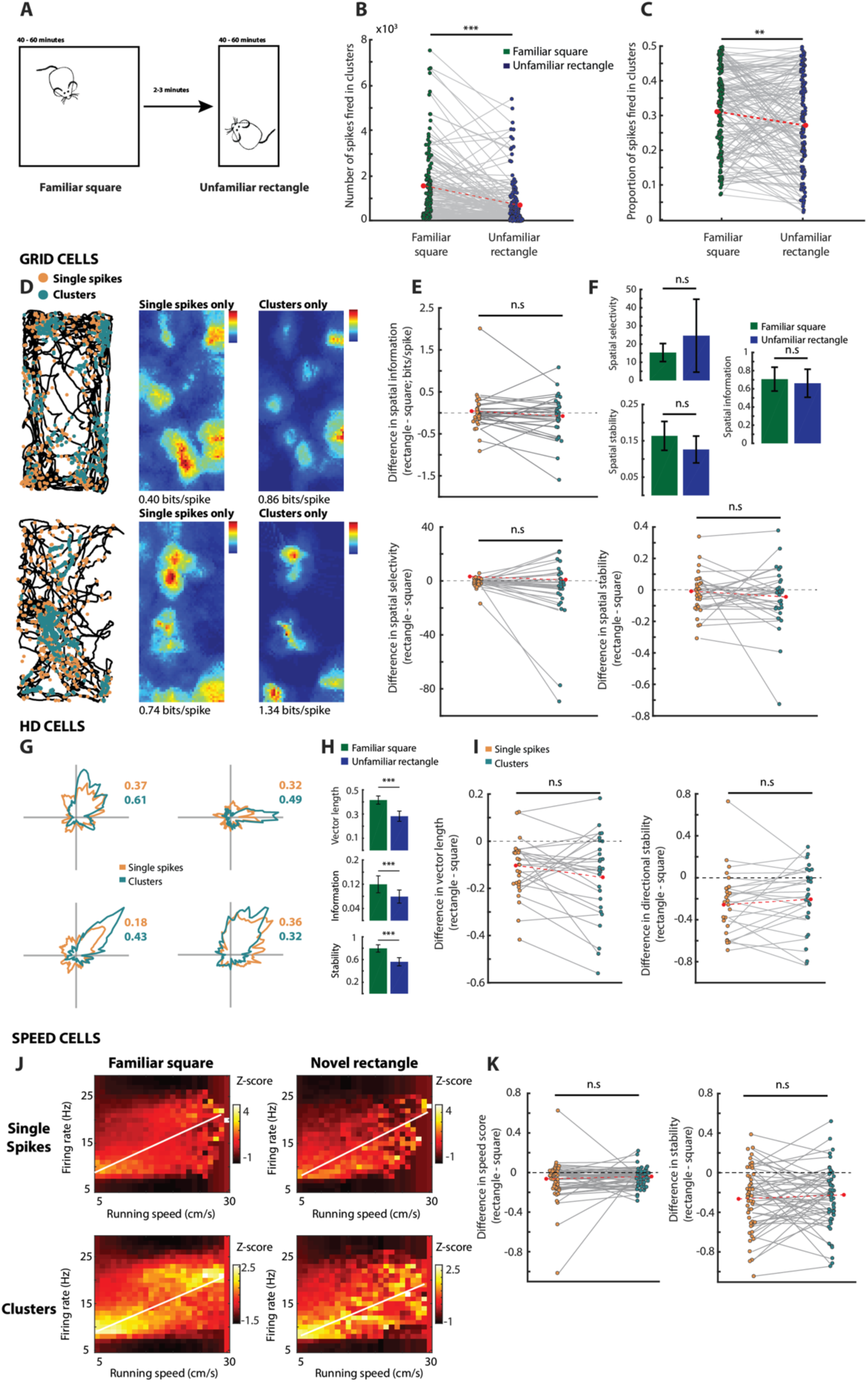
Environmental novelty reduces cluster proportion and changes cell coding properties. **A)** Schematic illustration of the recording procedure; Animals were initially recorded in a familiar square 1 m x 1 m box for 40-60 minutes (until their coverage of the envrironment was greater than 70%). If cells of interest were recorded, animals were removed from the square and transferred to a novel rectangular environment (0.5 m x 1 m) where they were recorded for the same approximate duration (with the same coverage criterion). **B)**. Cells fired significantly fewer of their spikes as clusters in the unfamiliar rectangle (purple circles) compared to the familiar square (green circles). This absolute reduction in the number of cluster spikes was representative of an overall lower proportion of total spikes fired as clusters **(C). D)** Two example grid cells recorded in the modified rectangular environment. Maps showing the path of the animal over recordings as black lines (left plots) are overlaid with the position that single spikes (orange) and cluster spikes (green) were fired. Rate maps generated from only the single spikes (center plots) show generally less compact fields with more outfield firing than the more precise rate maps generated from the burst spikes (rightmost plots). **E)** There was no overall difference in spatial information (top; Z = 1.10, p = 0.273, n.s), selectivity (middle; Z = 1.88, p = 0.060, n.s) or stability (bottom, 1.92, p = 0.055, n.s) between the familiar and unfamiliar environments, although information and stability trended toward lower scores in the unfamiliar rectangle. **F)** When broken down into individual comparisons in the difference in scores between environments for single spikes alone and clusters alone, there was no difference in the information rate (top left; Z = 1.25, p = 0.21, n.s), spatial selectivity (bottom left; Z = 1.63, p = 0.104, n.s) or spatial stability (bottom right; Z = 0.49, p = 0.624, n.s) between single spikes (orange) and clusters (green). **G)** Four individual head direction cells recorded in the unfamiliar rectangle, divided into the directional rate maps derived from their single spikes (orange) and their clusters (green) in isolation. The mean vector length produced from each map is shown color-coded alongside each map. In the unfamiliar environment clusters still tended to be more directionally precise than single spikes. **H)** Compared to their activity in the familiar square, there was a robust decrease in the mean vector length (top), directional information content (middle) and directional stability (bottom) when the same cells were recorded in the unfamiliar rectangle. On the other hand, there was no overall difference in the amount of change in any of these metrics between the single spikes and clusters of the cells; both decreased an equivalent amount (**I). J)** Z-scored probability plots of two speed cells in the familiar square (left column) and novel rectangle (right column). The single spikes fired by these cells are shown on the top row, while the clusters are illustrated on the bottom row. White lines on these plots show the slope and intercept of linear regressions through the firing rate/speed data. **K)** While there was a general decrease in speed score (left) and stability (right) between the environments, there was no difference in the change in either score or stability between clusters and single spikes. All figures show the zero point as a black dotted line, and the mean of all groups is shown with a red point connected by a dashed red line. All figures; n.s = not significant, * p < 0.05, ** p < 0.01, *** p < 0.001.

### Single spikes are more robustly entrained to theta than clusters

The majority of cells in MEC prefer to fire spikes toward the peak of LFP theta^32^, and the phase at which cells fire on the theta wave is thought to segregate the flow of information into distinct functional epochs^33^. We therefore examined the phase relationship of clusters of spikes compared to single spikes on the MEC theta wave (Figure 7A-C). Across all cells in MEC, there was a marked tendency for cells to fire toward the peak of LFP theta (Figure 7D). However, when comparing single spikes to clusters, single spikes were much more robustly entrained to theta than clusters. Cells identified as grid cells tended to fire at the trough of LFP theta (Figure 7E), and the single spikes of these cells were significantly more entrained to theta than the clusters (Figure 7F-G, mean phase vector, single spikes: 0.244 ± 0.031, clusters: 0.178 ± 0.030; Z = 3.018, p = 0.0025). Head direction cells were found to fire at either the peak or the trough of theta (Figure 7H,I); but again, the single spikes were robustly more phase locked than the clusters Figure 7J, mean phase vector, single spikes: 0.250 ± 0.028, clusters: 0.209 ± 0.025; Z = 3.661, p = 2.517^-4^). Speed cells likewise were entrained to the peak or trough of theta (Figure 7K), but like grid and head direction cells, the single spikes fired by these cells were robustly locked to theta, while the clusters were less so (Figure 7L,M), mean phase vector, single spikes: 0.227 ± 0.022, clusters: 0.191 ± 0.019; Z = 3.094, p = 0.0020).

**Figure 7.**
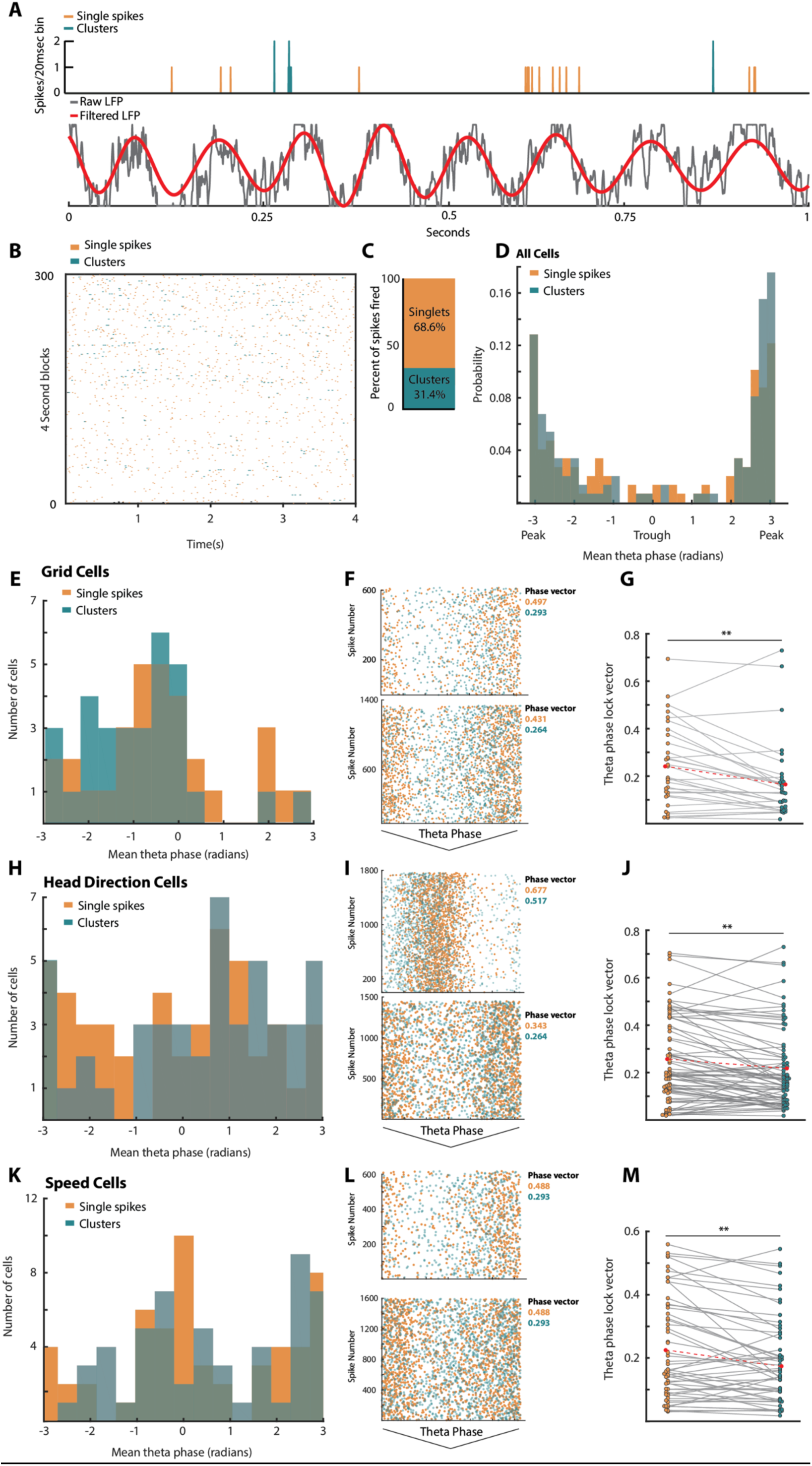
Single spikes are more strongly locked to LFP theta than clusters. **A)** Example 1 second recording showing the co-recorded spiketrain of a single cell (top) and LFP (bottom) illustrating the relationship of single spikes (orange) and clusters (green) to the LFP. Raw LFP is shown in grey, with bandpass-filtered LFP in red. **B)** Raster plot of the firing activity of a cell recorded over 1200 seconds, divided into 4-second blocks. Single spikes are shown in orange, with clusters shown in green. **C)** Stacked bar chart showing the relative proportions of spikes fired singly (orange) compared to in clusters of two or more (green) for the whole population of cells. **D)** The probability that a cell had a specific mean phasic directional relationship to theta for the entire population of cells. Both single spikes (orange) and clusters (green) of all cells tended to have a mean direction aligned to the peak of theta. **E)** As in (D), but for cells identified as grid cells. **F)** Example raster plots of two grid cells relative to the underlying theta phase divided into single spikes (orange) and clusters (green). Both cells are phase locked to the peak of theta, but the single spikes are more tightly phase locked than the clusters. The phase-locking vector of the single spikes and clusters are shown to the right of each plot. **G)** The phase-lock vector of each grid cell’s single spikes (orange) and clusters (green). Single spikes are significantly more phase locked to theta than clusters. Grey lines connect the divided spiketrains of each cell. The mean of each group is shown with a red dot and connected with a dashed red line**. (H-J)** as in (E-G), except for cells identified as directional. (H-I) Directional cells were divided between preferring the trough or peak of theta. (J) As with grid cells, single spikes fired by directional cells were significantly more locked to theta than clusters. **(K-M)** As in E-G and H-J, except for cells identified as speed cells. As with directional cells, speed cells tended to prefer to fire either at the peak or trough of theta, but the single spikes fired by these cells were significantly less locked to the local theta phase than the clusters of these cells.*p < 0.05, ** p < 0.01, *** p < 0.001.

## Discussion

As demonstrated previously, we find that neurons within MEC can be clustered into “bursty” neurons that fire many spikes within 10 ms of a previous spike, and non-bursty neurons that seldom did. Projection onto principal component space allowed successful k-means differentiation of these two groups. Moving beyond the traditional view of bursts as related spikes occurring within 10ms to a more liberal definition of “clusters”, we then subdivided spikes fired by a neuron into single, isolated spikes and clusters of spikes. When the spiketrains of neurons were divided into clusters and single spikes, a clear pattern emerged; the clusters fired by spatial and directional cells contained significantly more precise information than the equivalent number of single spikes. Cells identified as coding for speed also fired clusters of spikes that were better and more stably correlated with running speed that the equivalent number of singly-fired spikes. These observations are a direct measure that supports previous data suggesting bursts are correlated with more stable and precise spatial information^16,17^, and demonstrates that a clear delineation exists between clusters of spikes organized on the timescale of theta oscillations, and spikes fired at longer time intervals.

Since traditional bursts are associated with strong upstream input^13,14^ it is likely that clusters of spikes also tend to be fired when a variable-coding neuron within MEC receives information through a strongly potentiated pathway, or from several upstream sensory neurons simultaneously. Since it has been suggested through in-silico modelling^34^ and experimental evidence that neurons coding for different variables within MEC interact with one another, it is also possible that clusters are fired when this activity converges onto the cluster-firing neuron from many different intracortical connections. Whatever the case, the richer coding of space and direction by clusters suggests that these spikes are more computationally useful and meaningful, which we suggest represents a “certainty” mechanism.

In addition to the richer coding of space and direction by clusters, when animals were exposed to environmental uncertainty by exposure to a novel environment, all cell types fired fewer clusters overall. Crucially, they also fired fewer clusters as a proportion of their total number of spikes. This finding further supports the concept that clusters represent a “certainty” mechanism. The mechanism may be either through amplification of familiar sensory inputs, increased reciprocal feedback from hippocampus or another, less transparent mechanism. It is clear, however, that neurons fire proportionally more clusters in spatially familiar environments, and when environmental ambiguity is introduced the input to these neurons through well-established or potentiated connections may decrease, causing a larger proportion of isolated spikes to occur relative to clusters. This raises the possibility that the function of single spikes might be to “check” for changes in spatial geometry, so that cells can rapidly shift synaptic weights or connections to respond to novel environments.

One critical aspect of the present study is the suggestion that single spikes and clusters represent computationally distinct units. In support of this notion, we find that single spikes, while representing less well defined spatial, directional, and speed information than clusters of spikes, are much more strongly phase-locked to the LFP theta wave, while clusters are less phasically predictable but relay much less noisy information about space. This differentiation in the spatial and temporal accuracy of spikes and clusters suggests that they may represent computationally distinct modes of activity. Strong entrainment to local LFP would suggest coordination among many local neurons, while activity that is less locally phase-locked but more spatially “accurate” may represent a firing mode communicating information inter-regionally. Thus we propose that the precise coding of space and direction in the MEC is carried by a minority of spikes that are fired in clusters and which convey rigid, well-defined information about space. By contrast, the majority of spikes that are fired singly are less spatially precise, but are strongly entrained to theta enable “fuzziness” and rapid communication and adaption of inputs and outputs to and from MEC.

We therefore propose distinct functions for clusters versus single spikes; Single spikes attend closely to LFP theta and are more likely to operate at times synchronized between regions. Clusters, being less entrained to LFP theta are less likely to convey long-range information and are more likely to represent convergence of spatial information from multiple sources intraregionally. These findings have important implications for our understanding of connectivity both within MEC and between MEC and regions such as HPC. This differentiation between activity “modes” also potentially enriches computational models of the MEC/HPC loop.

## Materials and Methods

### Subjects

All procedures were carried out in accordance with and under the approval of the Institutional Animal Care and Use Committee at the Stanford University School of Medicine. Male C57/BL6 mice (n = 25) mice were initially housed in groups of between one and five animals, prior to surgery with food and water *ad libitum*. After animals received their surgical implants they were singly housed. Animals continued to have free access to water, but were food deprived such that they were maintained at 80% of their free-feeding weight. Animal housing was kept on a 12 h light/dark cycle, with all surgical and experimental procedures performed during the animal’s light phase. Animals weighed between 18-31 g at the time of surgery.

### Surgery

Buprenorphine (0.1mg/kg, i.p) was delivered to each animal immediately prior to surgery as a prophylactic analgesic. Animals were then induced to anaesthesia through a mixture of oxygen and isoflurane (2-3 %). Once animals were anaesthetized, they were transferred to a surgical frame where they were maintained on isoflurane anaesthesia (1.5 – 2 %). Animals were implanted unilaterally in the left hemisphere with 16 – channel tetrode-carrying microdrives (Axona Inc.) aimed toward the medial portion of the entorhinal cortex. The tetrodes themselves were composed of polyimide-coated alloy (90 % Platinum, 10% Iridium) wires that were 17 micron in diameter. Just prior to implantation, these electrodes were cut flat, and electroplated using Platinum Black (Neuralynx Inc) solution until their impedance was measured in the range 150-300 kW. The electrode arrays were implanted at a depth of 900-1000 microns, 3.25-3.3 mm lateral from the midline and 0.5mm rostral from the posterior sinus, at an approximate - 4° angle from the vertical plane. Microelectrodes were affixed to screws inserted into to the skull during surgery using dental acrylic cement. One of the screws had a ground wire soldered to it, which was soldered to the ground pin of the Microdrive. Post-surgery animals recovered on a heated pad (∼35° C) for several hours until fully ambulatory. Animals recovered for 3 days before habituation to the recording environment and 7d before recordings began. If necessary, during this recovery period animals were administered analgesia in the form of i.p injection of Carprofen. Physiological 0.9% saline was given subcutaneously during surgery and during recovery as needed to maintain adequate hydration.

### In vivo recordings

Animals were screened daily for the presence of single units by connecting them to the recording apparatus, which consisted of a dual-LED unity-gain op-amp headstage connected to digital preamplifier and a system control unit, which connected to a PC running system control software (DACQUSB, Axona Inc) and allowing them to freely explore a square 1 x 1 m environment until their path had covered at least 75 % of it. This typically meant that animals explored the square environment for between 30-60 minutes per day. Signals were digitally amplified between 3,000 – 10,000 times and then bandpass filtered between 800 and 6,700 Hz. LFP was acquired by digitally mirroring one of the channels and then lowpass filtering it 500 Hz (notch filter at 60 Hz). This signal was stored in high resolution at a 4.8kHz sampling rate and in lower resolution at a sampling rate of 250 Hz. The dual-LED assembly in the headstage allowed tracking and storage of the animal’s position in space. Left was distinguished from right due to the size of the LED assemblies (the left side was marked by four individual LEDs in a square configuration, while the right was marked by two LEDs in a line). The entire recording apparatus was surrounded by a removable black curtain which reached nearly to the roof of the room. The enclosure was dimly lit by a 100W lamp outside the curtain which reflected from the corner of the ceiling.

At the conclusion of the day’s recording, tetrodes were lowered ∼25 mm by turning a screw on the Microdrive approximately 1/8 of a full turn (45°). If, during a day’s recording a cell of interest was identified, the animal was removed from the recording chamber, the chamber was cleaned through application of Nature’s Miracle odor-neutralizing spray, and then a removable wall was placed into the environment halfway along one axis, producing a 1 m x 0.5 m rectangular box. The animal was returned to this recording environment with novel geometry, and recorded for a further 20-60 minutes (depending on rate of coverage). After this “modified environment” recording, the animal’s electrodes were lowered by ∼25 mm as in the case in which no cells of interest were found.

### Histology

On completion of experiments animals were deeply anaesthetised with sodium pentobarbitone and then perfused transcardially with 0.9 % saline followed by 10 % paraformaldehyde. Animals were decapitated, the brains removed and stored in 10 % formalin for 24 h, and then transferred to 30 % sucrose solution until the brains sank to the bottom of the container. Brains were then sectioned sagittally into 40 micron thick slices, mounted, and then stained with cresyl violet. Electrode position was determined microscopically by examining each slide. Experiments were considered “completed” when either the animals had exposure to the novel rectangular environment 13 times, or the recordings became excessively noisy, indicating the electrodes had left the brain, or several days passed without any new cells being recorded, whichever criterion was reached first.

### Spike-sorting and downsampling

After recordings and offline, spikes were classified manually into clusters using vendor-specific cluster-cutting software (TINT, Axona, Inc.). Clustering was performed by comparing various aspects of the waveform feature space on each electrode of a tetrode. Initial clustering was done using amplitude, and then refined using other waveform features such as time of peak to ensure good isolation of cells. Cluster separation and quality was determined post-cutting by computing the distance in Mahalanobis space between cells on the same tetrode (as in Schmitzer-Tobert et. al^35^. If a spike was either preceded by or followed by a spike within 100 ms, it was classified as a cluster spike, otherwise it was classified as a single spike.

Many of the spatial variables compared in the present study can be affected by the number of spikes. In order to control for differences in the number of spikes classified as clusters or single spikes, we performed downsampling on the spike trains. Whichever part of the spiketrain (single spikes or clusters) contained a greater number of spikes was randomly downsampled such that it matched the length of the shorter part. This was done by randomly selecting spikes from the longer spiketrain (without replacement) and deleting them.

### Cell-type Classification

Cells were initially classified according to their tuning curves compared against a shuffled distribution as has been done previously^36^. (Speed cell criteria, correlation > 0.05, stability > 0.4; Head direction cell criterion, mean vector length > 0.2; Grid cell criterion, grid score > 0.3). Grid scores were calculated from the correlation of the grid maps at 60° and 120° minus the correlations at 30°, 90° and 150°. Head direction scores were the mean directional vector computed from the directional angle of the head of the animal at which spikes were fired. Speed scores arise from the correlation between firing rate and running speed, and speed stability was calculated from the mean correlation of the speed/firing rate relationship over four temporal bins spanning the recording.

### Analysis of EEG

EEG was recorded simultaneously with single units, and recordings were therefore included only under the same criteria as single units (i.e. > 70 % coverage). Data was sampled at either 250 Hz or 4.8 kHz, or both. The highest sampling rate trace was used where available. In order to perform analysis of theta, EEG was bandpass filtered between 5 and 11 Hz, and a power spectrum was derived using a Fourier transform. A Hilbert transform was performed on filtered data to derive instantaneous power, phase, and frequency.

### Statistics

All statistical tests were performed in MATLAB (2017b) and were two-sided. All data was tested for parametricity using a Lilliefors test before determining whether to use parametric or nonparametric comparisons. Circular mean directions, vector length calculations, and statistical comparisons were performed using the CircStat package for MATLAB^37^. In all cases where data did not conform to a normal distribution, Wilcoxon signed-rank tests were used for paired data, and rank-sum tests were used for unpaired data.

## Acknowledgments

RGK was funded by the Neurological Foundation of New Zealand while the data in this manuscript were collected. These data were collected in the laboratory of Lisa Giocomo at Stanford University and some data previously appeared in Munn et. al. (2020).

## Author Contributions

R.G.M Conceptualized the experiments and analyses and collected data. L.R.Q and S.M.T provided support on analyses. R.G.M wrote the paper with feedback and input from L.R.Q and S.M.T.

